# Effects of UniProtKB restructuring and taxonomic database restrictions on downstream Unipept peptide-centric profiling

**DOI:** 10.64898/2026.02.24.707692

**Authors:** Tibo Vande Moortele, Simon Van de Vyver, Ben-Björn Binke, Tim Van Den Bossche, Lennart Martens, Peter Dawyndt, Pieter Verschaffelt, Bart Mesuere

## Abstract

Metaproteomics identifies the proteins present in a microbial community and, from them, which organisms are active and what functions they carry out. Tools such as Unipept assign peptides to taxa and functions by matching them to UniProtKB proteins and taking the lowest common ancestor of the matching taxa, deriving functional annotations from the same matches. This analysis therefore depends directly on the content of UniProtKB, which was substantially restructured in 2025–2026, reducing it by more than 100 million protein entries. We investigated how these changes affect taxonomic interpretation and how restricting the database to sample-specific taxa identified by rRNA profiling modifies that effect. Fixed, non-redundant peptide lists from human-gut and marine-hatchery studies were reanalysed with Unipept against UniProtKB releases 2025_03, 2025_04, and 2026_02. Peptide mapping coverage fell from 85.9% to 73.3% (gut) and from 82.3% to 68.7% (marine) between release 2025_03 and 2026_02, yet dominant taxonomic profiles remained robust: family- and genus-level distributions changed little and all 15 dominant gut species were retained. Root-level assignments dropped from 18.7% to 7.0% (gut) and 25.8% to 14.3% (marine). Restricting UniProtKB 2026_02 to taxa detected by large-subunit ribosomal RNA (LSU rRNA) profiling reduced coverage to 64.4% (gut) and 41.8% (marine), while genus- and species-level proportions changed by less than one percentage point. The major UniProtKB restructuring of 2025-2026 therefore narrowed mapping breadth without destabilizing the dominant taxonomic profiles in these two datasets.

**Significance:** UniProtKB underpins many database-dependent metaproteomics workflows. How it is curated quietly shapes which organisms are detected, and how confidently they can be named. Its 2025-2026 restructuring is the resource’s largest change in years, removing around 142 million of its 253 million entries and adding 39 million new ones. Yet how such release-to-release changes propagate into downstream peptide-to-taxon annotation has not been systematically examined, nor whether analyses run against one release remain trustworthy once it changes. Using Unipept and two contrasting datasets as a test case, this work addresses two questions: (i) how successive UniProtKB releases affect the reproducibility of peptide-centric taxonomic annotation, and (ii) the value and risks of tailoring the reference database with external metagenomic evidence. These lessons are practical. While the dominant taxonomic results hold across UniProtKB versions, meaningful differences remain. This highlights how strongly the underlying reference database shapes metaproteomic analyses and reconfirms the importance of accurately recording which database version was used.

## Introduction

Microbial communities drive many processes ranging from biogeochemical cycles to host health. Characterizing them requires knowing not only which organisms are present but which are metabolically active, and which functions they carry out. Meta-omics approaches address complementary aspects of this question. Metagenomics describes the genes a community could express, while metatranscriptomics describes the genes that are transcribed. Metaproteomics, by contrast, measures the proteins actually present. This is the most direct readout of the realized function and of the taxa contributing it [1–4]. By identifying peptides through mass spectrometry and mapping them to proteins and taxa, metaproteomics links community composition to metabolic activity across diverse environments, including the human gut [5–7] and marine [8,9] ecosystems. This peptide-to-protein-to-taxon mapping depends directly on the reference protein database: a peptide can be identified and assigned a taxon only when a matching protein is present. An incomplete or taxonomically skewed reference can leave peptides unmatched, shift their assignments to less informative ranks, or bias the inferred community profile. The quality and composition of the reference protein database are therefore a fundamental determinant of metaproteomic accuracy.

In database-search workflows, tandem mass spectra are first matched to peptide sequences; those identified peptides can then be mapped to reference proteins and taxa for downstream interpretation. Reference-database composition can affect both stages, but through different mechanisms. At the spectrum-to-peptide stage, database size and composition affect computational burden and statistical error control [10]. At the downstream annotation stage studied here, Unipept remaps already identified peptide sequences to matching proteins and taxa. Here, taxonomic breadth matters more than raw entry count: matches spanning distant or weakly specified taxa can move a peptide’s lowest common ancestor (LCA) [11] upward, whereas removing such matches can produce a more specific LCA or eliminate the match.

Database selection for spectrum-to-peptide identification has been studied extensively [12–14], and tools such as Unipept Desktop [15] support targeted reference-database construction. Less is known about how release-to-release changes in UniProtKB propagate into downstream peptide-to-taxon annotation when the identified peptide set is held constant.

Unipept [11,15–17] is a peptide-centric analysis platform that provides taxonomic and functional views of metaproteomic peptide sets. Unlike protein-inference workflows that aggregate peptide evidence into protein groups, Unipept treats each peptide independently: it identifies matching UniProtKB [18] proteins and assigns the lowest common ancestor of their associated taxa. Alterations in database content and taxonomic representation can therefore change whether a peptide maps, the rank and identity of its LCA, and the aggregate taxonomic peptide-support profile [19].

Some UniProtKB entries are linked to non-standard or weakly informative NCBI taxonomy [20] nodes, including “unclassified,” “uncultured,” and environmental lineages. When a peptide matches both a well-defined lineage and one of these distant or weakly specified branches, its LCA may move to a less informative rank, including the root. Unipept has therefore historically applied a taxon filter that prunes designated problematic branches before LCA calculation [11].

In 2025 and 2026, UniProt underwent a staged restructuring of its protein space. Release 2025_04 introduced a new reference-proteome selection and contained both removals and additions. Release 2026_02, released on June 10, 2026, completed the transition to a UniProtKB composed of reference-proteome entries, reviewed records, and selected unreviewed entries with experimental or other biologically important evidence [18]. The net result was a decrease from approximately 254 million to 149.8 million entries, accompanied by substantial changes in sequence and taxonomic representation. These changes are relevant to every UniProtKB-dependent workflow because they alter which sequences and taxa are available and because analyses performed against different releases may not be directly reproducible. The public releases also include annotation and taxonomy updates; comparisons among them therefore capture the net effect of restructuring rather than the isolated effect of database size.

Auxiliary taxonomic profiles can guide construction of smaller protein reference databases [21–23]. Rather than using the auxiliary profile to identify peptides, taxa supported by that profile determine which protein entries remain available for downstream peptide-to-taxon mapping. Such restriction can remove competing matches and produce more-specific LCAs, but it can also make peptides unmatched or redistribute their LCAs when the auxiliary profile omits relevant taxa or domains or when taxonomy conversion is imperfect.

In this study, the auxiliary evidence comprised large-subunit ribosomal RNA (LSU rRNA) taxonomic profiles available through study-linked MGnify analyses for the gut and marine studies [25]. Using the family-level taxa these profiles support, we restricted the UniProtKB 2026_02 database to gut- and marine-specific subsets; we refer to the comparison of each restricted subset with unrestricted 2026_02 as LSU-informed restriction. This restriction changes only which protein entries are available for downstream mapping and does not modify the fixed peptide lists.

Despite the broad importance of UniProtKB, the downstream consequences of its recent restructuring for peptide-centric taxonomic annotation remain unclear. Using Unipept as a focused use case, we held two previously identified peptide sets fixed and ask three questions: (1) how do selected UniProtKB releases spanning the restructuring affect peptide mapping coverage, LCA-rank distribution, root-level assignments, and taxonomic assignment patterns; (2) how does LSU-informed restriction of UniProtKB 2026_02 alter mapping coverage, conditional LCA resolution, root assignments, and taxon composition; and (3) how does the incremental effect of Unipept’s taxon filter change across the selected releases? This design isolates downstream annotation effects and does not repeat the original spectrum-to-peptide database searches.

## Materials and methods

### Study design and scope

This study evaluates three factors in a common downstream Unipept workflow: selected UniProtKB release, dataset-specific LSU-informed database restriction, and Unipept’s taxon filter. The same fixed, non-redundant peptide list for each dataset was used in every comparison, and the original spectrum-to-peptide searches were not repeated.

First, the peptide lists were mapped against selected UniProtKB releases 2025_03, 2025_04, and 2026_02 with Unipept’s taxon filter enabled to measure release-associated changes in mapping coverage, LCA-rank distribution, root-level assignments, and taxonomic assignment patterns. Second, 2026_02 was restricted to families reported by the corresponding LSU analysis and compared with unrestricted 2026_02, with the Unipept’s taxon filter enabled in both configurations. Third, the three selected global releases were analysed with Unipept’s taxon filter enabled and disabled to quantify its incremental effect; this on/off comparison was not applied to the LSU-restricted databases. The workflow is summarized schematically in **Figure 1**.

**Figure 1:**
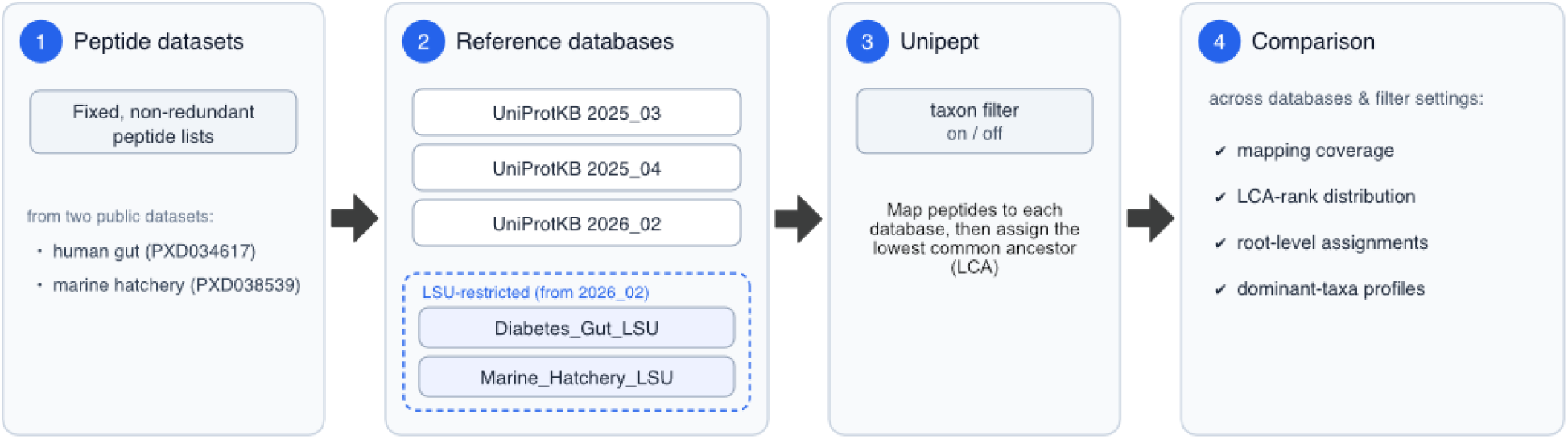
Overview of the study workflow. 1) Two public metaproteomics datasets (human gut and marine hatchery) were preprocessed into fixed, non-redundant peptide lists. 2) Reference databases were assembled: three successive UniProtKB releases (2025_03, 2025_04, and 2026_02) and, for each dataset, an LSU-informed subset of the 2026_02 release built from MGnify LSU rRNA taxonomic profiles. 3) Each peptide list was annotated with Unipept against every database using the lowest-common-ancestor approach, with Unipept’s taxon filter enabled and disabled. 4) The resulting taxonomic assignments were compared across databases and filter settings in terms of mapping coverage, LCA-rank distribution, root-level assignments, and dominant-taxa profiles.

### Metaproteomics datasets

Peptide data were derived from two publicly available metaproteomics datasets obtained from the PRIDE repository [24] under accessions PXD034617 and PXD038539 (**Table 1**), which were reported by Wang et al. [25]. These data are a reanalysis of the data originally published by Heintz-Buschart et al. [26] and Timmins-Schiffman et al. [27]. These datasets respectively correspond to the gut and marine hatchery microbiome. The human gut dataset was collected from 18 individuals from four families, with at least two members in each family having type 1 diabetes mellitus. This resulted in a total of 36 mzIdentML files [28]. The marine hatchery dataset was collected from twelve water samples across three time points (36 mzIdentML files).

**Table 1:** Summary of the metaproteomics dataset used as input for Unipept analyses.

| Feature | PXD034617 (Human Gut) | PXD038539 (Marine Hatchery) |
| --- | --- | --- |
| **Number of peptides** | 317 136 | 68 603 |
| **Number of unique peptides** | 67 799 | 8 182 |
| **Original Study** | PXD003791 (Type 1 Diabetes) | PXD020692 (Marine Microbiome) |
| **Environment** | Human Gut (Host + Microbes) | Marine / Shellfish Microbiome |
| **Search DB** | MGnify Assemblies + UniProt | MGnify Assemblies + UniProt |
| **Search Engines** | X!Tandem [29,30] & MS-GF+ [31] | X!Tandem & MS-GF+ |
| **Peptide Length** | 5 – 50 amino acids | 6 – 60 amino acids |
| **Precursor Tol.** | 10 ppm | 20 ppm |
| **Fragment Tol.** | 0.02 Da | 0.02 Da |
| **Identified Proteins** | 59 613 | 268 |

Peptides were extracted using a custom XML parser (the Python script can be found in the supplementary material). All peptides passing the FDR threshold were retained. Prior to downstream analysis, peptide sequences were deduplicated to generate non-redundant peptide lists for Unipept. For the human gut, only the 20 mzIdentML files linked to diabetes were retained.

For the human gut, a total of 317136 peptides were found, corresponding to 67 799 unique peptide sequences. For the marine hatchery dataset, a total of 68603 peptides were found, corresponding to 8182 unique peptides. These datasets were used consistently across all database configurations and filtering strategies evaluated in this study. **Table 1** provides a detailed summary of both datasets.

### Reference protein databases

To characterize release-associated changes in downstream annotation, we compared three selected UniProtKB releases spanning the 2025–2026 restructuring (**Table 2**):

1. UniProtKB release 2025_03 (Jun 18, 2025), hereafter the baseline: the last release before the restructuring, containing approximately 254 million protein sequences.
2. UniProtKB release 2025_04 (Oct 15, 2025), hereafter the intermediate release: the initial large-scale restructuring, which removed redundant proteins and proteins from taxonomically unclassified organisms, leaving approximately 199 million proteins.
3. UniProtKB release 2026_02 (Jun 10, 2026), hereafter the latest release: the final result of UniProt’s restructuring and curation, containing approximately 150 million protein sequences.

**Table 2:** List of databases. The first three rows are different releases of UniProtKB. The last two rows represent databases that are constructed by restricting UniProtKB 2026_02 to families derived from the corresponding LSU rRNA taxonomic profiles.

| Database Name | UniProt Version | # Proteins | # Taxa |
| --- | --- | --- | --- |
| UniProtKB_2025_03 | 2025_03 | ~ 254 million | ~ 1.3 million |
| UniProtKB_2025_04 | 2025_04 | ~ 199 million | ~ 1.3 million |
| UniProtKB_2026_02 | 2026_02 | ~ 150 million | ~ 219 thousand |
| Diabetes_Gut_LSU | 2026_02 | ~ 41 million | ~ 19 thousand |
| Marine_Hatchery_LSU | 2026_02 | ~ 23 million | ~ 11 thousand |

Release contents and restructuring details were obtained from UniProt’s online documentation (https://www.uniprot.org/help/refprot_only_changes).

In addition, LSU-informed family restrictions were constructed using large subunit (LSU) ribosomal RNA-based taxonomic identifications from MGnify [32]. For the human gut and marine hatchery dataset, accessions MGYS00001985 and MGYS00005863 respectively contain the corresponding taxonomic assignments. These Genome Taxonomy Database (GTDB [33]) notations were converted to NCBI taxonomic identifiers at the family level. Mapping was performed at the family level because GTDB-to-NCBI correspondence is most reliable at higher ranks; species- and genus-level GTDB lineages frequently lack a one-to-one NCBI equivalent, which would drop legitimate taxa from the filter.

These NCBI taxonomic identifiers were used to construct custom taxon-restricted databases in Unipept. For each dataset, we generated a database based on their LSU identifications. **Table 2** summarizes the resulting databases.

### Unipept configuration and annotation workflow

All analyses were performed using the Unipept web application (version 6.4.3; https://unipept.ugent.be), accessed on 8 July 2026. The isoleucine/leucine (I/L) equivalence rule was enabled for all analyses. As of version 6.0.0, missed cleavage handling is enabled by default in Unipept [17]. All analyses handle tryptic, semi-tryptic and non-tryptic peptides, as well as peptides that contain missed cleavages.

Taxonomic annotation was performed using Unipept’s LCA approach. Each peptide was assigned to the most specific taxonomic level shared by all matching proteins. Rank-level percentages are reported cumulatively: a peptide resolved to a given rank is also counted at every less specific rank on the same lineage to the root of the tree, so the family-level value includes all peptides assigned at family level or a more specific rank (genus or species). These rank percentages are therefore nested rather than mutually exclusive and are not expected to sum to 100%. All LCAs were then aggregated into comprehensive community profiles that were used to compare all database configurations.

Throughout this study we distinguish five quantities. First, mapping coverage is the number of unique input peptides with at least one protein match, expressed as a percentage of all unique input peptides. Second, cumulative LCA-rank yield is the number of peptides resolved to a given rank or a more specific one, expressed as a percentage of all unique input peptides. These nested values are the rank percentages reported in the main text. Third, conditional LCA-rank yield is that same count expressed as a percentage of matched peptides rather than of all input peptides. Fourth, the Sankey diagrams show mutually exclusive exact-LCA categories. Each peptide contributes to exactly one node (root, domain, class, order, family, genus, species, or unmatched), the node values use all input peptides as the denominator, and they sum to 100% only when the unmatched category is included; they are therefore not cumulative. Fifth, the top-taxa bar charts show each taxon’s share of the species-resolved peptide evidence, defined as the number of peptides whose exact LCA is that species divided by the total number of species-resolved peptides. The “Other” category aggregates all species-resolved peptides outside the top 15. Because this normalization is taken within a species-resolved set whose size changes between databases, absolute supporting-peptide counts are reported in the text so that compositional shifts cannot conceal changes in the underlying peptide evidence.

Unipept’s taxonomic filter [11] was evaluated as part of the study design. This filter is baked into Unipept and removes unclassified or unwanted taxa according to rules defined in Unipept, pruning problematic branches before the LCA is computed. Analyses were performed both with and without this filter to assess its necessity and impact in the context of successive database restructuring. Unless stated otherwise, all analyses reported in the main text and figures use Unipept’s taxon filter.

## Results

We first quantified how the successive UniProtKB releases affected downstream peptide mapping and taxonomic assignment, then how LSU-informed database restriction changed these outcomes, and finally the incremental contribution of Unipept’s taxon filter. Each analysis was carried out for both the human-gut and marine-hatchery datasets, and the following sections report them in turn.

### Effects of successive UniProtKB restructuring

To assess the effect of different UniProtKB databases on peptide-centric metaproteomics analysis, we compared the baseline (2025_03), intermediate (2025_04), and latest (2026_02) releases. Analyses were performed independently for both the gut microbiome and marine hatchery datasets.

#### Mapping coverage across selected releases

For the gut microbiome dataset (67 799 unique peptides), 58 222 peptides (85.9%) were matched using the baseline. Following the initial large-scale restructure in the intermediate release, 52 730 peptides (77.8%) retained at least one protein match, and in the latest release 49 720 peptides (73.3%) were matched. The marine hatchery dataset (8 182 unique peptides) followed the same downward trend: 6 735 peptides (82.3%) were matched using the baseline, decreasing to 6 042 peptides (73.8%) in the intermediate release and 5 617 peptides (68.7%) in the latest release.

From 2025_03 to 2026_02, mapping coverage declined by 12.5 percentage points in the gut dataset and 13.7 points in the marine dataset. Nevertheless, 73.3% of gut peptides and 68.7% of marine peptides remained mappable with the latest selected global release despite a net reduction of more than 100 million UniProtKB entries. This demonstrates substantial downstream mapping resilience alongside a meaningful loss of mapping breadth.

#### Cumulative LCA-rank yields across selected releases

In the gut microbiome dataset, family-, genus-, and species-level assignments using the baseline respectively accounted for 42.8%, 34.4% and 23.2% of total unique peptides on a cumulative basis (each rank includes all peptides assigned at that rank or a more specific one) (**Figure 2**). Following the initial restructuring, family-level assignments increased slightly to 44.0%, while genus- and species-level assignments decreased to 33.2% and 20.3%. Using the latest release, 42.5% were classified at the family level, 31.5% at the genus level, and 19.8% at the species level. Thus, successive database restructurings were accompanied by a marginal redistribution toward higher-rank LCA assignments and a slight reduction in species-level resolution (23.2% to 19.8%).

**Figure 2:**
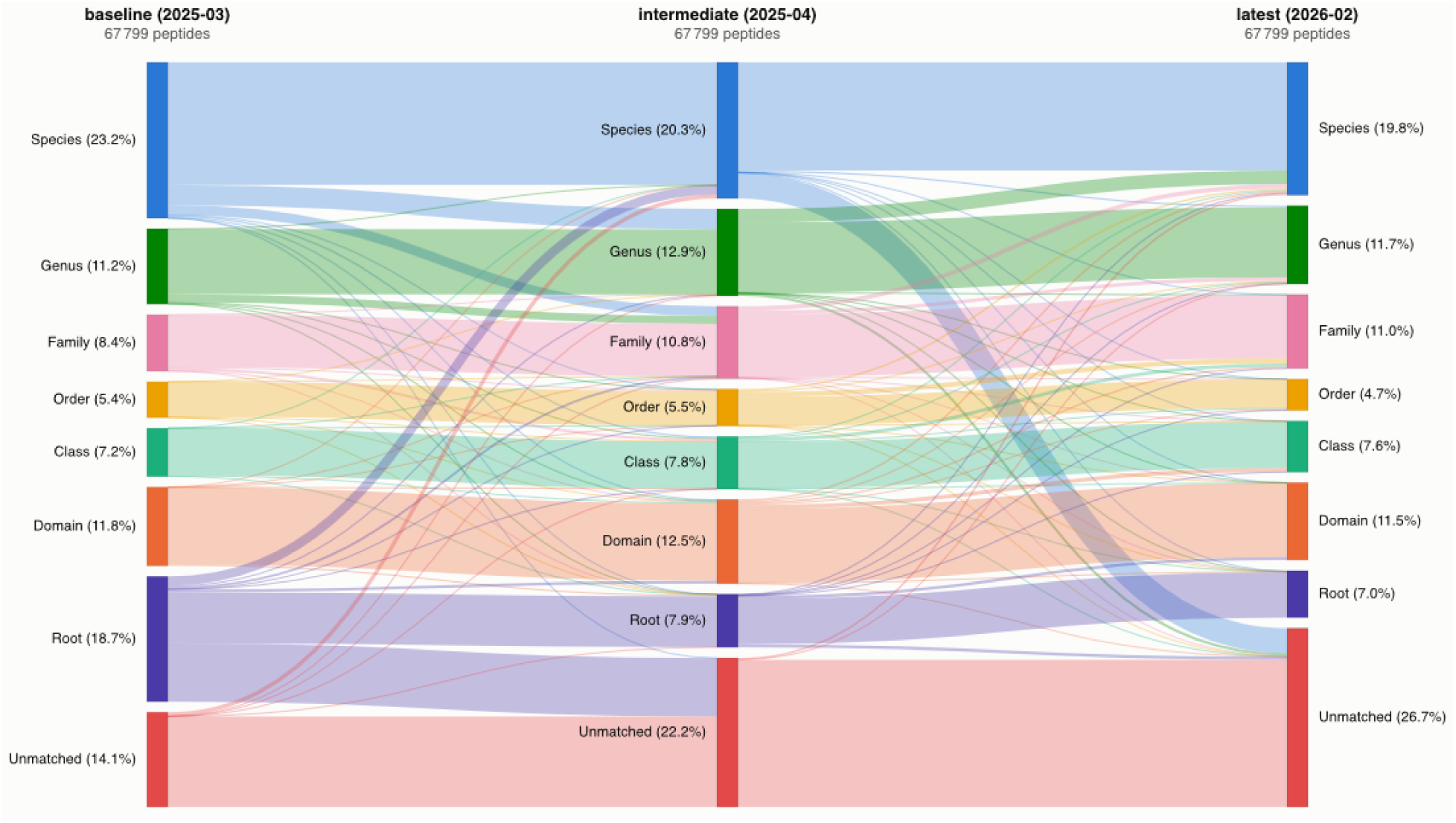
Sankey diagram showing peptide-level transitions among exact LCA-rank categories for the human gut dataset across three successive UniProtKB releases. The three columns correspond, from left to right, to UniProtKB 2025_03 (baseline), 2025_04 (intermediate), and 2026_02 (latest). Flows between adjacent columns show how each peptide’s exact LCA changed from one release to the next. Each node is a mutually exclusive exact-LCA category (root, domain, class, order, family, genus, species, or unmatched). Node values are shares of all 67 799 input unique peptides and sum to 100% only when the unmatched category is included, so they are not cumulative.

In the marine dataset, family-, genus-, and species-level assignments using the baseline accounted for 19.7%, 14.2%, and 12.0% of peptides, respectively. These remained highly stable in the intermediate release (19.7%, 14.1%, 11.8%) and the latest release (19.2%, 14.2%, 12.1%). In contrast to the gut dataset, the marine dataset was more resilient, with taxonomic resolution remaining nearly constant despite the substantial decrease in underlying protein search space (**Supplementary, Figure 1**).

Conditional on being matched, these release-associated changes appear as coverage effects rather than losses of resolving power. In the gut microbiome dataset the species-resolved fraction of matched peptides remained essentially constant across releases (27.0%, 26.1%, and 27.0% for 2025_03, 2025_04, and 2026_02), even though the per-input species-level yield fell from 23.2% to 19.8%; the decline therefore tracks the loss of mapping coverage rather than a reduced ability to resolve matched peptides to species. In the marine hatchery dataset the conditional species-resolved fraction instead increased across releases (14.6%, 16.0%, and 17.6%), consistent with a stable per-input yield over a shrinking matched set.

#### Reduction in root-level ambiguity and peptide transitions

Root-level assignments decreased substantially in both datasets. In the gut dataset, the root share fell from 18.7% with 2025_03 to 7.9% with 2025_04 and 7.0% with 2026_02. In the marine dataset, it fell from 25.8% to 18.1% and 14.3%, respectively. The baseline-to-latest reductions were therefore 11.7 percentage points in gut and 11.5 points in marine, directly demonstrating fewer least-informative LCA outcomes.

The peptide-level transitions in **Figure 2** and **Figure S1** resolve this root decline into three outcomes for the root-assigned peptides in the baseline: they either lost their matches and became unmatched, remained at the root, or gained a more specific rank. In the gut dataset, 48% of the baseline-root peptides became unmatched, 35% remained at the root, and 17% gained a more specific rank. In the marine dataset, 43% became unmatched, 50% remained at the root, and 7% gained a more specific rank. Because the releases combine entry removals and additions with annotation and taxonomy changes, the aggregate root decline should not be equated with improved taxonomic accuracy or attributed to a single mechanism.

#### Preservation of dominant taxonomic profiles

To assess whether successive UniProtKB restructuring changed Unipept’s inferred taxonomic community composition, we compared profiles of species-resolved peptide evidence across database configurations. Analyses focused on the top 15 taxa by peptide evidence in the baseline and evaluated their presence and relative ordering in the other two configurations.

For the gut microbiome dataset, the top 15 species by peptide evidence, identified using the baseline, were also found using the intermediate and latest releases (**Figure 3**). The same pattern was observed at the family and genus levels. No dominant species disappeared entirely following the transition from the baseline to newer configurations. Although higher-rank taxonomic profiles were stable, notable differences were observed for certain species-level assignments in the gut microbiome dataset.

**Figure 3:**
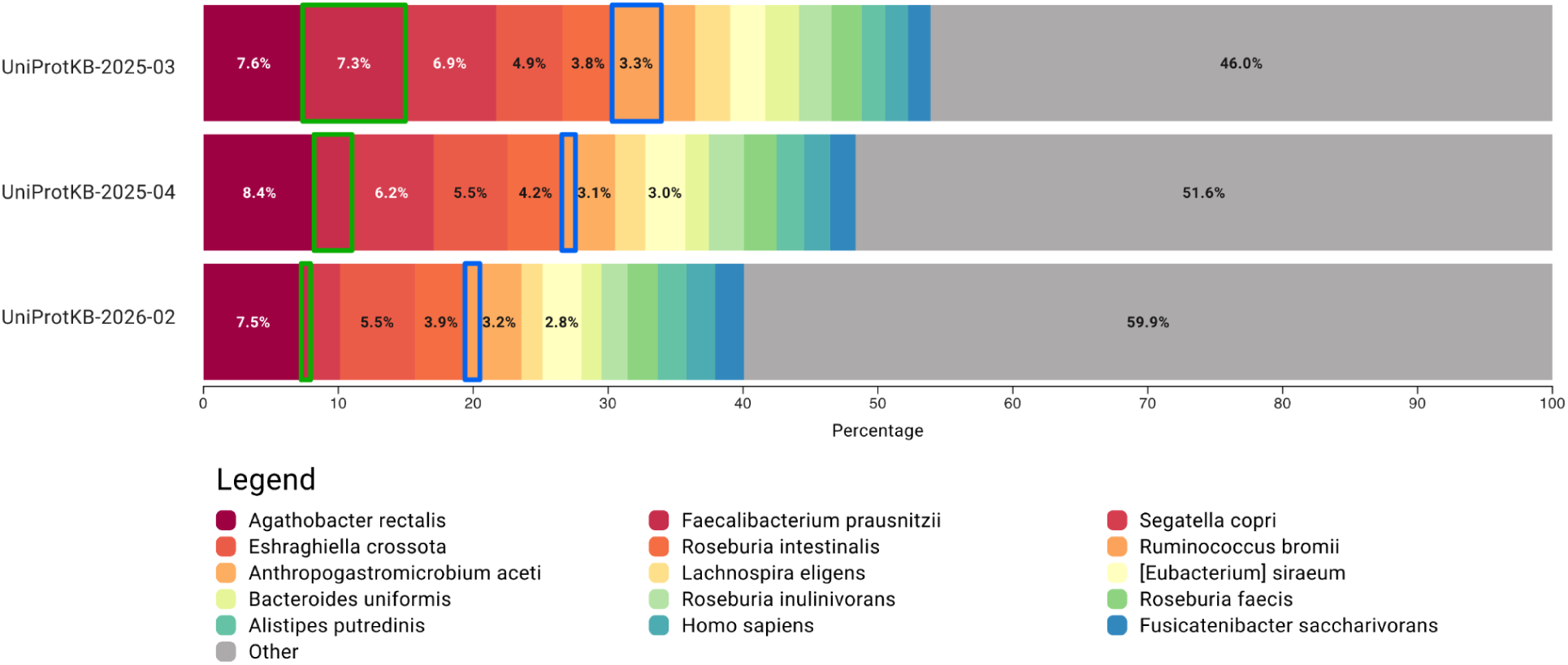
Top 15 gut species ranked by peptide evidence using the UniProtKB 2025_03 configuration as a baseline. Bars show each species’ share of the species-resolved peptide evidence, and “Other” aggregates all species-resolved peptides outside the top 15. All taxa found in UniProtKB 2025_03 are also picked up by the new, reduced UniProtKB configurations. *Faecalibacterium prausnitzii* (green border) and *Ruminococcus bromii* (blue border), however, show a marked relative reduction and are highlighted in the three different configurations.

In particular, *Faecalibacterium prausnitzii* showed reduced peptide evidence across UniProtKB configurations. At the genus-level, however, peptide evidence for *Faecalibacterium* remained the same across all database versions. Further inspection indicated that peptides previously assigned specifically to *F. prausnitzii* in the baseline were redistributed across closely related species and reference proteomes in the reduced databases. This explains the stability at the genus rank.

Similarly, *Ruminococcus bromii* exhibited a substantial reduction in species-level peptide evidence for the intermediate and latest releases. This corresponded to a considerable reduction in the number of protein entries associated with this species between database versions (approximately 192 000 proteins in the baseline versus 115 000 in the latest release).

For the marine hatchery dataset, taxonomic profiles were generally stable across database configurations at all three levels (family, genus, and species). The top 15 taxa by peptide evidence, identified in the baseline, were all recovered using the intermediate release with highly similar peptide evidence (**Figure 4**).

**Figure 4:**
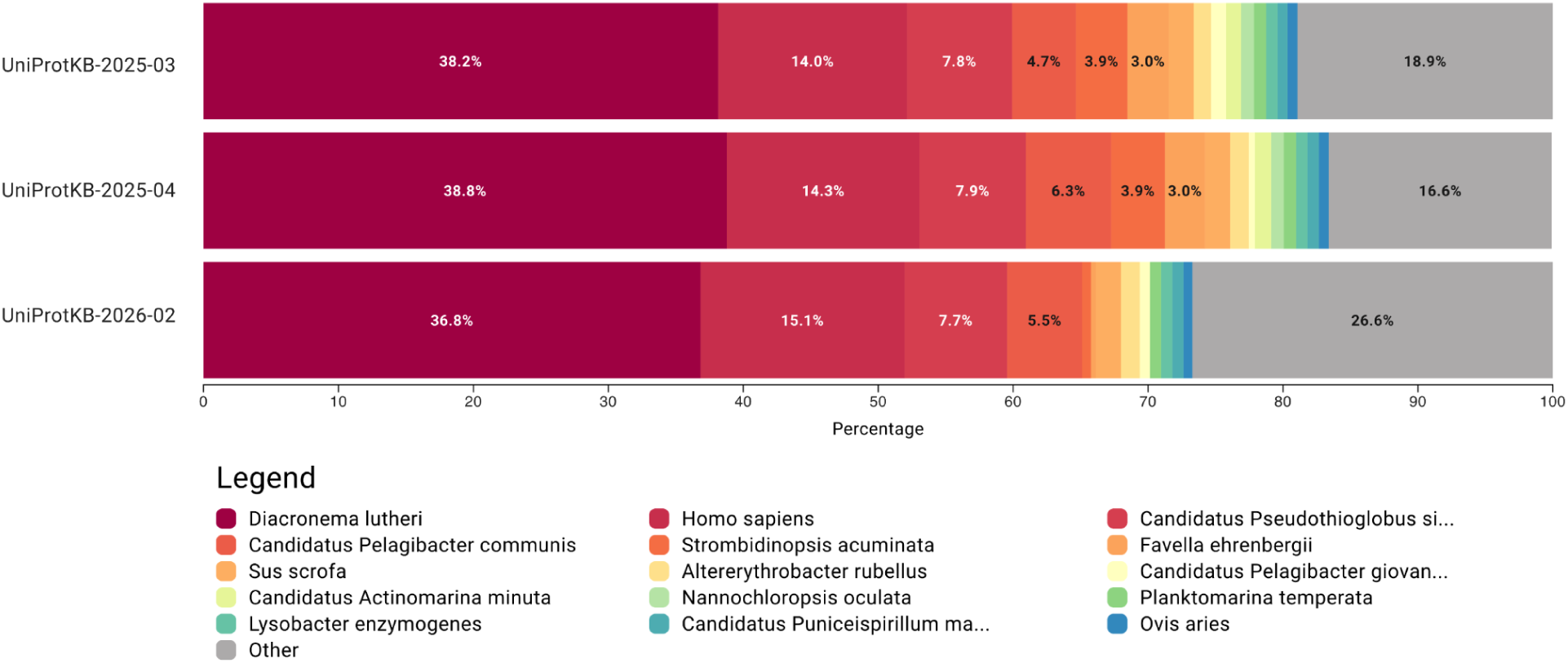
Top 15 marine hatchery species ranked by peptide evidence using the UniProtKB 2025_03 configuration as baseline. Bars show each species’ share of the species-resolved peptide evidence, and “Other” aggregates all species-resolved peptides outside the top 15. *Candidatus Actinomarina minuta* and *Nannochloropsis oculata* were both absent for the most recent configuration due to the lack of a corresponding reference proteome.

However, in the latest release, certain family-, genus-, and species-level taxa were no longer detected. These taxa corresponded to organisms for which no reference proteome was available in UniProtKB at the time of analysis (e.g. *Candidatus Actinomarina minuta* and *Nannochloropsis oculata*). Consequently, peptides previously assigned to these taxa could not be matched by the more restricted latest release.

### Impact of LSU-informed restriction

For this analysis we restricted the latest release (2026_02) to the families detected in each environment by their LSU rRNA profiles, producing environment-specific databases for the human gut and marine hatchery that we compared against the unrestricted 2026_02 release. The following sections report the resulting effects on mapping coverage, conditional taxonomic resolution, root-level assignments, and dominant taxa.

#### Mapping coverage across LSU-restricted databases

For the gut microbiome dataset (67 799 unique peptides), 49 720 peptides (73.3%) were matched using the unrestricted latest release (2026_02), and LSU-informed restriction reduced this to 43 664 peptides (64.4%). The effect was more pronounced in the marine hatchery dataset (8 182 unique peptides), where the unrestricted latest release matched 5 617 peptides (68.7%) and the LSU-restricted database matched only 3 424 peptides (41.8%). LSU-informed restriction substantially reduced peptide coverage in both environments.

#### Cumulative LCA-rank yields across LSU-restricted databases

Expressed as a percentage of all unique input peptides, family-, genus-, and species-level assignments accounted for 42.5%, 31.5%, and 19.8% under the unrestricted latest release and 43.3%, 31.5%, and 19.7% under the LSU-restricted database for the gut microbiome dataset (**Figure 5**). The absolute number of species-level assignments per input peptide was therefore essentially unchanged (13 425 versus 13 343 peptides). For the marine hatchery dataset the corresponding per-input values were 19.2%, 14.2%, and 12.1% (unrestricted) versus 28.6%, 13.6%, and 11.1% (LSU-restricted) (**Supplementary, Figure S2**). Species-level assignments fell from 991 to 911, a difference of only 0.98 percentage points.

**Figure 5:**
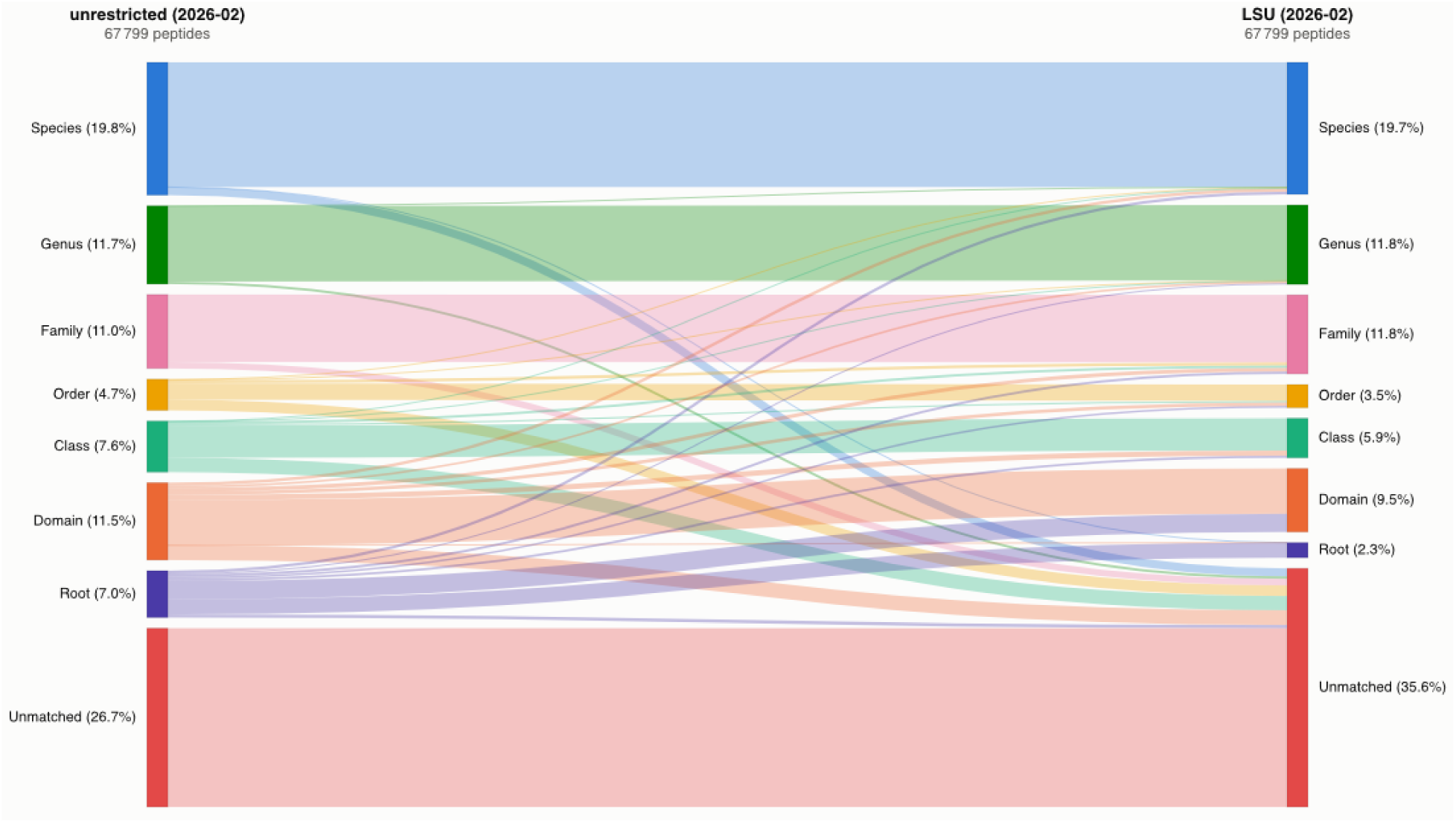
Sankey diagram showing human gut peptide assignments between two UniProtKB 2026_02 configurations. The two columns correspond, from left to right, to the unrestricted 2026_02 release (unrestricted) and the LSU-restricted database (LSU). Flows between the columns show how each peptide’s exact LCA changed under restriction. Each node is a mutually exclusive exact-LCA category (root, domain, class, order, family, genus, species, or unmatched). Node values are shares of all 67 799 input unique peptides and sum to 100% only when the unmatched category is included, so they are not cumulative.

Conditional on being matched, the contrast is stronger because the matched set itself shrank. In the gut microbiome dataset the species-resolved fraction of matched peptides rose from 27.0% (13425 of 49 720) to 30.6% (13343 of 43 664), and in the marine hatchery dataset from 17.6% to 26.6%. Further, the family-level fraction of matched peptides rose from 27.9% to 68.3% in the marine dataset. LSU restriction thus nearly preserved the absolute number of species-level assignments per input peptide while markedly increasing the species-resolved fraction of the much smaller matched set.

Across both datasets, LSU-informed restriction reduced the matched set more than it reduced the number of species-level assignments, so species-level assignments accounted for a larger fraction of the matched peptides. This shift was small in the gut dataset, but more pronounced in the marine dataset.

#### Reduction in root-level ambiguity and peptide transitions

For the gut microbiome dataset, 7.0% of peptides were assigned to the taxonomic root using the latest release, and the LSU-restricted database further reduced this to 2.3%. The marine hatchery dataset followed the same pattern, with root-level assignments decreasing from 14.3% in the latest release to 5.1% under the LSU-restricted database.

The peptide-level transitions in **Figure 5** and **Figure S2** resolve this root decline into three outcomes for the peptides assigned to the root under the unrestricted latest release: they either lost their only match and became unmatched, remained at the root, or gained a more specific rank. In the gut dataset, 7% of these root peptides became unmatched, 32% remained at the root, and 61% gained a more specific rank. In the marine dataset, 9% became unmatched, 36% remained at the root, and 55% gained a more specific rank. Unlike the release restructuring, LSU restriction therefore lowered root ambiguity mainly by reassigning root peptides rather than dropping them. These reassignments were concentrated at higher ranks, however, with most reaching only the domain-to-family levels and few the genus or species level (8% in the gut dataset and 10% in the marine dataset).

#### Preservation of dominant taxonomic profiles

To determine whether metagenomics-derived filtering altered inferred community composition, we compared the share of species-resolved peptide evidence using the LSU-restricted database to the profile derived from the unrestricted latest release. Because these profiles are normalized within the species-resolved set, whose size differs between configurations, we report them alongside absolute supporting-peptide counts (**Figures 6** and **7**) so that compositional shifts are not mistaken for changes in absolute support. In the marine dataset, for example, the top 15 species by peptide evidence accounted for 526 of 991 species-resolved peptides (53%) under the unrestricted release but 726 of 911 (80%) under the LSU-restricted database. This apparent concentration reflects both genuine gains for some taxa (*Diacronema lutheri*, 365 to 420 supporting peptides) and the appearance of low-support or biologically implausible taxa admitted by the family-level restriction (*Neotoma lepida*, 0 to 18; *Mesocricetus auratus*, 0 to 12), rather than a uniform rise in peptide evidence.

**Figure 6:**
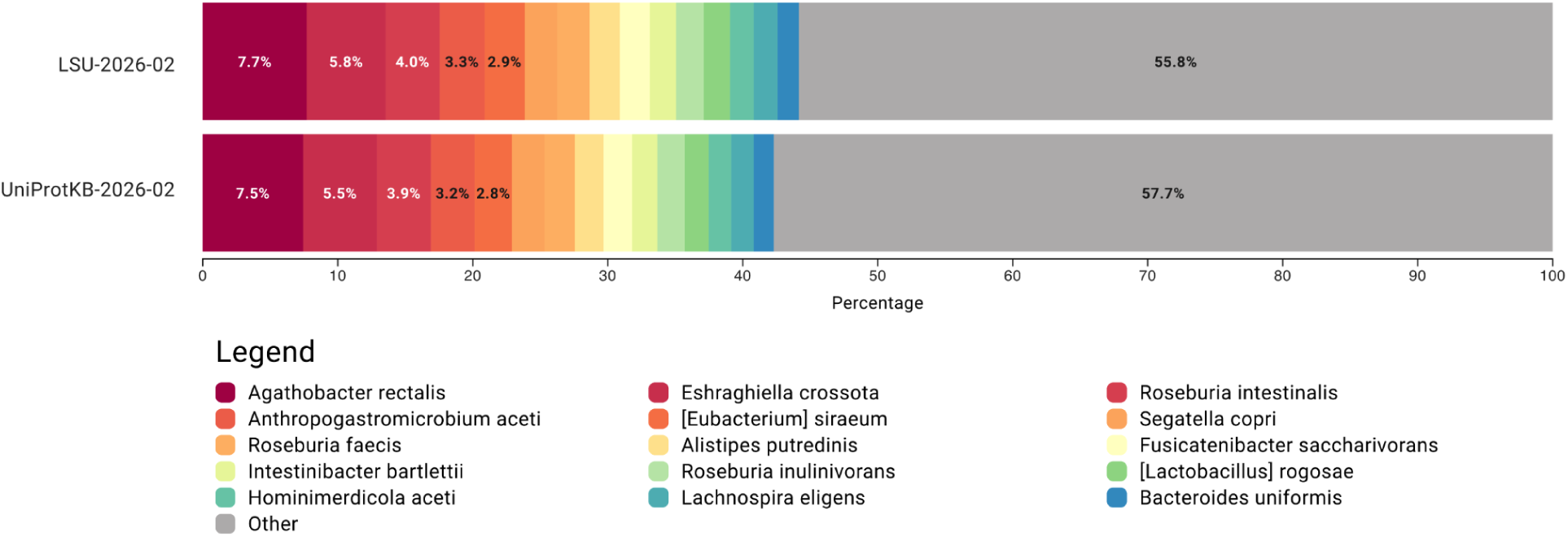
Top 15 gut species ranked by peptide evidence under the LSU-restricted database, compared against the unrestricted 2026_02 release. Bars show each species’ share of the species-resolved peptide evidence. The “Other” category aggregates all species-resolved peptides outside the top 15.

**Figure 7:**
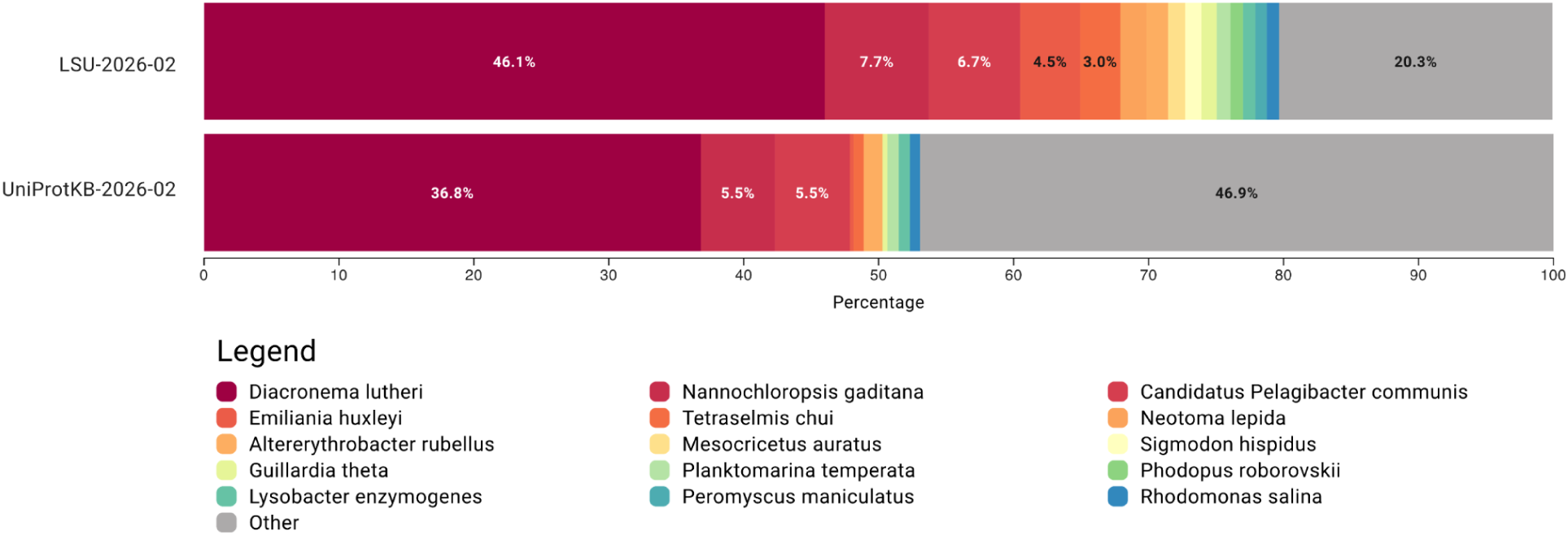
Top 15 marine hatchery species ranked by peptide evidence under the LSU-restricted database, compared against the unrestricted 2026_02 release. Bars show each species’ share of the species-resolved peptide evidence. The “Other” category aggregates all species-resolved peptides outside the top 15. Substantial differences in both resolution and support were seen across the two configurations.

For the gut microbiome dataset, the top 15 species by peptide evidence were similar under the LSU-restricted database (**Figure 6**). The same pattern was observed at the family- and genus-level. Peptide evidence remained highly comparable, and no substantial shifts in evidence occurred among these dominant taxa. For example, the top 15 species accounted for 5 683 of 13 425 species-resolved peptides (42%) under the unrestricted release and 5 883 of 13 343 (44%) under the LSU-restricted database, with individual supporting-peptide counts near-identical (e.g. *Agathobacter rectalis*, 1 001 versus 1 032; *Roseburia intestinalis*, 529 versus 536). One notable difference was the presence of *Homo sapiens* in the unrestricted configuration, which was absent in the LSU-filtered analyses, consistent with its exclusion from the LSU-derived family list.

Despite the overall stability of dominant taxa, some species were exclusively detected in either the unrestricted or filtered databases. Nine species (with at least 10 supporting peptides) were exclusively observed using the unrestricted database. Five of those are linked to the human gut microbiome (*Agathobaculum hominis, Anaerotignum faecicola, Evtepia gabavorous, Flintibacter faecis* and *Gemmiger formicilis*). The other four species included *Gallus gallus, Candidatus Onthenecus intestinigallinarum, Homo sapiens* and *Sus scrofa*. Conversely, *Zea mays* was exclusively found by the metagenomics-derived filter. *Zea mays* is a terrestrial crop plant, so its exclusive detection by the LSU-restricted database most likely reflects an artifact of the family-level GTDB-to-NCBI mapping rather than a biologically meaningful gut taxon.

For the marine hatchery dataset, we observed substantial differences in the top 15 taxa by peptide evidence of the LSU-restricted database (**Figure 7**). These differences were visible at all three taxonomic levels (species, genus and family). Using the unrestricted UniProtKB database, almost all of the top 15 taxa by peptide evidence appeared either with substantially reduced peptide evidence or were not found at all.

Similar to the gut microbiome dataset, some species were detected only in the filtered or the unrestricted database. Five species (with at least 10 supporting peptides) were exclusively found using the unrestricted latest release. *Candidatus Pseudothioglobus singularis*, *Favella ehrenbergii* and *Strombidinopsis acuminata* are all found in marine contexts. The other two species were *Homo sapiens* and *Sus scrofa*, analogous to the gut microbiome dataset. Four species, with a minimum of 10 supporting peptides, were exclusively identified by the LSU-restricted database. All four (*Emiliania huxleyi*, *Mesocricetus auratus*, *Neotoma lepida* and *Scherffelia dubia*) were among its top 15 taxa by peptide evidence. Two of these, *Mesocricetus auratus* (golden hamster) and *Neotoma lepida* (desert woodrat), are terrestrial mammals whose appearance among the marine-hatchery taxa is biologically implausible. It probably reflects an artifact of the family-level GTDB-to-NCBI taxonomic mapping rather than genuine detection, and such hits should be screened out before biological interpretation.

This contrast suggests that the impact of LSU-informed restriction on taxon discoverability is heavily dataset-dependent. For the gut microbiome dataset, LSU-informed restriction did not reveal additional dominant taxa beyond those already detected using the unrestricted UniProtKB database, and overall taxonomic structure remained highly comparable. For the marine dataset, by contrast, targeted filtering detected certain taxa that were not observed using the unrestricted database. This indicates that database restriction can alter taxon discoverability in environments where coverage or taxonomic representation is more limited.

### Evaluation of Unipept’s taxon filter

To test whether Unipept’s taxon filter remains necessary as UniProtKB becomes more curated, we compared analyses with the filter enabled and disabled for each release, in both datasets.

#### Changes in taxonomic resolution

For the gut microbiome dataset, Unipept’s taxon filter substantially altered taxonomic resolution for the baseline (2025_03). Species-level assignments increased from 12.7% without filtering to 23.2% with filtering (**Figure 8**), with highly similar improvements at the genus and family levels. Using the intermediate release, this effect was smaller. Species-level assignments increased slightly from 19.0% to 20.3%, while genus- and family-level assignments showed only minor differences. For the latest release (2026_02), the impact of Unipept’s taxon filter was negligible, with differences of less than one percentage point at family, genus, and species levels.

**Figure 8:**
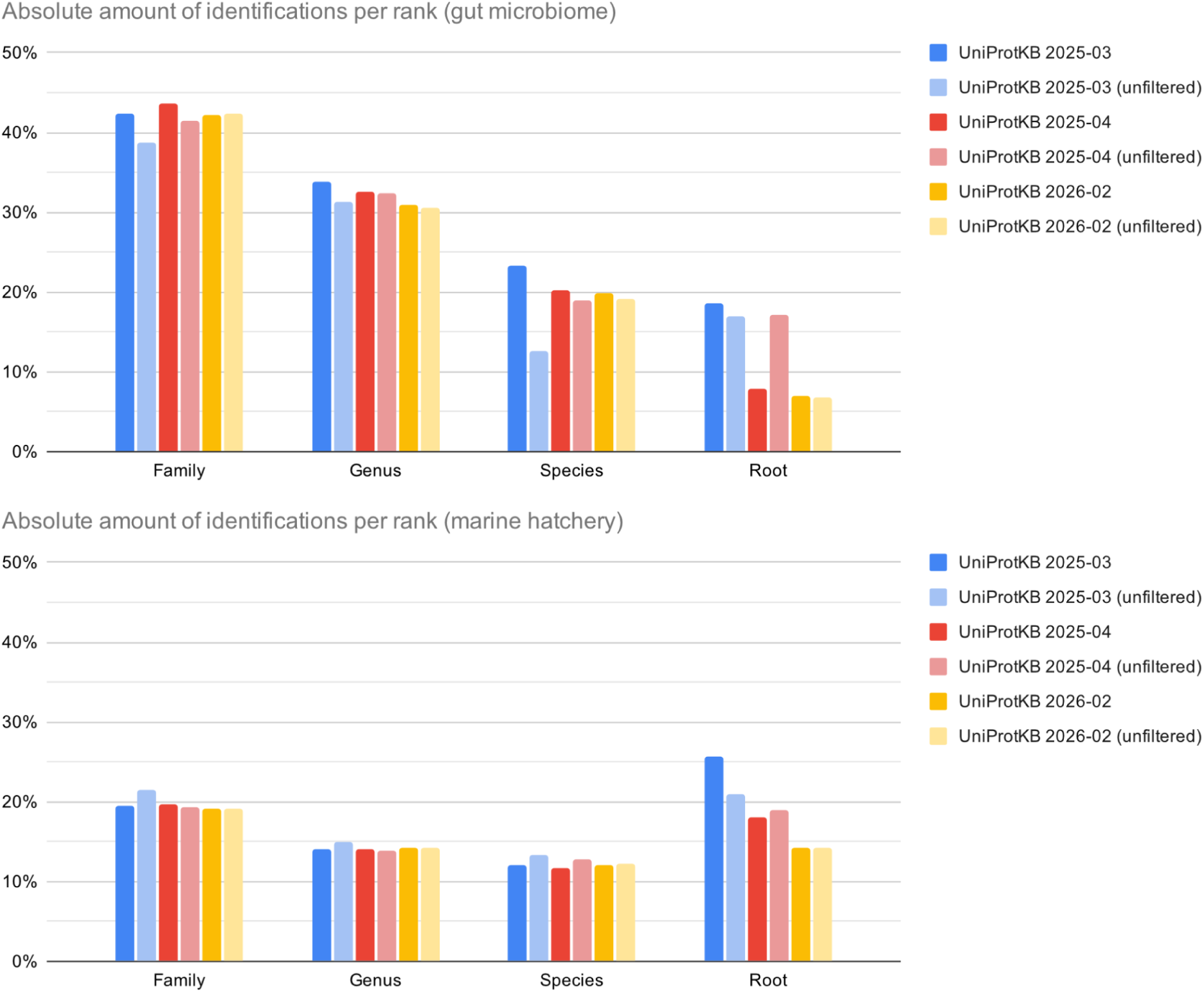
Comparison between three different configurations of UniProtKB (Unipept’s taxon filter enabled vs. disabled). Each chart shows the percentage of identifications per taxonomic rank (family-, genus- and species-level) on a cumulative basis. The top chart shows the results of the gut microbiome dataset. The bottom chart shows the identifications of the marine hatchery dataset. The percentage of peptides reduced to the taxonomic root is also shown for both datasets.

For the marine hatchery dataset, Unipept’s taxon filter did not improve lower-rank taxonomic resolution. Using the baseline, species-level assignments decreased slightly when filtering was applied (13.3% to 12.0%), with similar patterns at the genus and family levels, suggesting the filter may be too restrictive here. For the intermediate and latest releases, differences between analyses with and without the taxon filter were minimal (<1% absolute difference).

Overall, the impact of the taxon filter on taxonomic resolution varied between datasets and diminished from the baseline to the latest UniProtKB release, indicating effective curation by UniProt.

#### Reductions in root-level assignments

Unexpectedly, Unipept’s taxon filter did not always lower root-level assignments. For the gut microbiome dataset it raised them slightly in the baseline (17.0% without filtering to 18.7% with filtering), but lowered them in the newer releases. The reduction was largest in the intermediate release (16.2% to 7.9%), while in the latest release root-level assignments were already low (6.9%) and barely changed with filtering enabled (7.0%).

The marine dataset showed the same baseline reversal, more strongly: the filter increased root-level assignments from 20.9% to 25.8%. The effect shrank in the intermediate release (19.0% with the taxon filter disabled vs. 18.1% enabled) and disappeared in the latest release, where analyses with and without the taxon filter were nearly identical (14.3% vs. 14.2%).

## Discussion

Across two contrasting fixed peptide sets, the 2025–2026 UniProtKB restructuring narrowed downstream mapping breadth without destabilizing the dominant taxonomic signal inferred by Unipept. Most peptides remained mappable, family- and genus-level patterns were strongly conserved, all 15 dominant gut species persisted, and root-level assignments fell by more than 11 percentage points in both datasets. These findings provide direct evidence of robust downstream profiling rather than merely an absence of obvious failure. They do not, however, establish the biological accuracy of individual assignments or separate the effects of entry removal, entry addition, reference-proteome selection, annotation updates, and taxonomy updates.

Despite the removal of more than 100 million protein entries between the baseline and the latest release, Unipept’s taxonomic analyses remained robust. In both datasets, over two-thirds of unique peptides were still recoverable even for the 2026_02 release. Taxonomic profiles, particularly at the family and genus levels, were largely preserved. In the gut microbiome dataset, modest reductions in species-level assignments were observed. However, dominant taxa remained detectable and their relative ordering remained stable across releases. No previously dominant taxon disappeared in the gut dataset. The only losses occurred in the marine latest release, for organisms lacking a reference proteome (*Candidatus Actinomarina minuta*, *Nannochloropsis oculata*). Apparent shifts in peptide evidence for specific species such as *Faecalibacterium prausnitzii* and *Ruminococcus bromii* reflected redistribution of peptide assignments across closely related taxa and reference proteomes rather than disappearance of the broader genus-level assignment. In contrast, the marine dataset exhibited almost complete stability in taxonomic resolution across database versions, highlighting the resilience of LCA-based aggregation to large-scale database changes.

Successive database restructuring was accompanied by substantial decreases in root-level assignments, consistent with removed entries disproportionately contributing to ambiguous mappings. Reduced redundancy and the removal of taxonomically unclassified organisms both push root assignments down, and because UniProtKB applied them together, we cannot tell how much each one contributed. Regardless of the relative weighting, database downsizing shifted the balance toward more specific and less ambiguous assignments rather than representing a simple loss of resolution. These findings suggest that ongoing UniProtKB curation does not undermine metaproteomic workflows and may even enhance the resolution of taxonomic profiling.

Restricting UniProtKB using LSU-informed family lists introduced a clear trade-off between peptide coverage and conditional taxonomic resolution. In both datasets, targeted filtering substantially reduced the number of matched peptides while leaving the absolute number of species-level assignments per input peptide almost intact, so the retained species-level assignments made up a larger fraction of a smaller matched set, most markedly in the marine dataset. Restriction therefore did not so much create new species-level identifications as remove low-information matches around those already present. This also shows that reducing the search space alone does not guarantee more species-level identifications: even within a restricted database, peptides shared among closely related species continue to resolve to higher ranks. Thus, while LSU-informed restriction strongly reduced root-level assignments and concentrated resolution within the matched set, it did not increase the absolute number of species-level identifications.

The environment proved critical in determining the consequences of filtering. For the gut microbiome dataset, dominant taxa identified using the unrestricted database were largely preserved under restriction, and peptide evidence patterns remained highly comparable. In contrast, for the marine dataset, restriction substantially altered taxon discoverability and peptide evidence patterns. Several taxa identified as having strong evidence under the LSU-restricted database were either reduced or absent in the unrestricted configuration, and vice versa. This dataset-dependent contrast demonstrates that LSU-informed restriction can meaningfully alter taxonomic inference in environments where reference protein coverage is skewed or incomplete. While LSU-informed restriction can reduce ambiguity and remove implausible taxa (e.g., contaminants), it also carries the risk of excluding biologically relevant organisms if metagenomic detection is incomplete or subsequent taxonomic mapping is imperfect. The same imperfect, family-level mapping can also inject biologically implausible taxa (for example, terrestrial mammals in the marine filter and *Zea mays* in the gut filter), which should be identified and screened out before interpretation. Family-level mapping is therefore best understood as an explicit sensitivity-specificity trade-off rather than a default. It maximizes retention of related organisms at the cost of admitting broad or implausible hits. A finer-grained mapping would suppress these artifacts but risk discarding genuinely present taxa. Therefore, these LSU-informed database restriction strategies should be applied cautiously, considering environmental context and reference database completeness.

The impact of Unipept’s taxon filter decreased progressively across successive UniProtKB configurations. In the baseline it clearly improved species-level resolution in the gut dataset. These gains shrank in the intermediate release and vanished in the latest release. This is consistent with the filter mainly correcting for the problematic taxonomic assignments in UniProtKB that were more common in the older releases. Contrary to its intended effect, Unipept’s taxon filter raised root-level assignments in both baselines rather than lowering them. This occurs because pruning a branch is not the same as dropping a match. When a peptide’s only match sits on an invalid branch, the taxon filter prunes that branch and the peptide is reassigned to the root instead of being removed. Such invalid branches are rare in well-curated UniProtKB releases, so this case does not appear as often.

Overall, these findings indicate that as UniProtKB becomes more curated and centered on reference proteomes, the necessity of additional taxonomic filtering diminishes. This has practical implications for peptide-centric taxonomic analysis: reliance on aggressive taxon filtering may become less critical as UniProtKB evolves.

This study isolates downstream annotation by remapping two pooled, non-redundant peptide sets. It does not evaluate spectrum-to-peptide identification, sample-level variability, quantitative organism abundance, or taxonomic accuracy against ground truth. The UniProtKB release comparisons represent the net effect of simultaneous changes in entries, reference proteomes, annotations, and taxonomy. LSU-informed restriction additionally depends on the sample correspondence, aggregation rule, domain coverage, and completeness of the auxiliary taxonomic profile and on the accuracy of source-taxonomy-to-NCBI conversion. These boundaries limit generalization beyond the tested datasets, but do not alter the observed robustness of their dominant downstream profiles.

## Conclusion

In two fixed, non-redundant peptide datasets from human gut and marine environments, UniProtKB’s 2025–2026 restructuring reduced downstream Unipept mapping coverage by 12.5 percentage points in the gut dataset and 13.7 points in the marine dataset, yet dominant taxonomic profiles remained robust. Family- and genus-level patterns were strongly conserved, all 15 dominant gut species persisted, and root-level assignments fell by more than 11 points in both datasets. LSU-informed restriction imposed a pronounced dataset-dependent trade-off: it further reduced mapping coverage, increased the fraction of remaining matches assigned at more-specific LCA ranks, and substantially reorganized the dominant marine taxon profile. With UniProtKB 2026_02, the incremental effect of Unipept’s taxon filter was below one percentage point at every examined rank and at the root in both datasets. Thus, major UniProtKB restructuring narrowed mapping breadth without destabilizing dominant downstream Unipept profiles in these datasets. Effects on spectrum-to-peptide identification and other workflow stages were not tested.

## CRediT authorship contribution statement

**Tibo Vande Moortele:** Conceptualization, Data curation, Investigation, Methodology, Validation, Visualization, Writing – original draft, Writing – review and editing.

**Simon Van de Vyver:** Writing – review and editing.

**Ben-Björn Binke:** Writing – review and editing.

**Tim Van Den Bossche:** Writing – review and editing.

**Lennart Martens:** Writing – review and editing.

**Peter Dawyndt:** Writing – review and editing.

**Pieter Verschaffelt:** Conceptualization, Data curation, Methodology, Supervision, Writing – original draft, Writing – review and editing.

**Bart Mesuere:** Conceptualization, Supervision, Writing – review and editing.

## Ethics approval and consent to participate

Not applicable

## Data availability

No new mass spectrometry data were generated in this study. The metaproteomics datasets and the metagenomic taxonomic profiles reanalysed here are publicly available; their repositories and accession numbers are given in the Materials and Methods. The custom scripts used to parse the mzIdentML files and to construct the LSU-restricted database configurations, together with the resulting Unipept project files, are openly available at Zenodo (https://doi.org/10.5281/zenodo.22705564). All analyses were performed with the publicly available Unipept web application (https://unipept.ugent.be).

