## Supplementary for "Effects of UniProtKB restructuring and taxonomic database restrictions on downstream Unipept peptide-centric profiling"

### Supplementary Figures

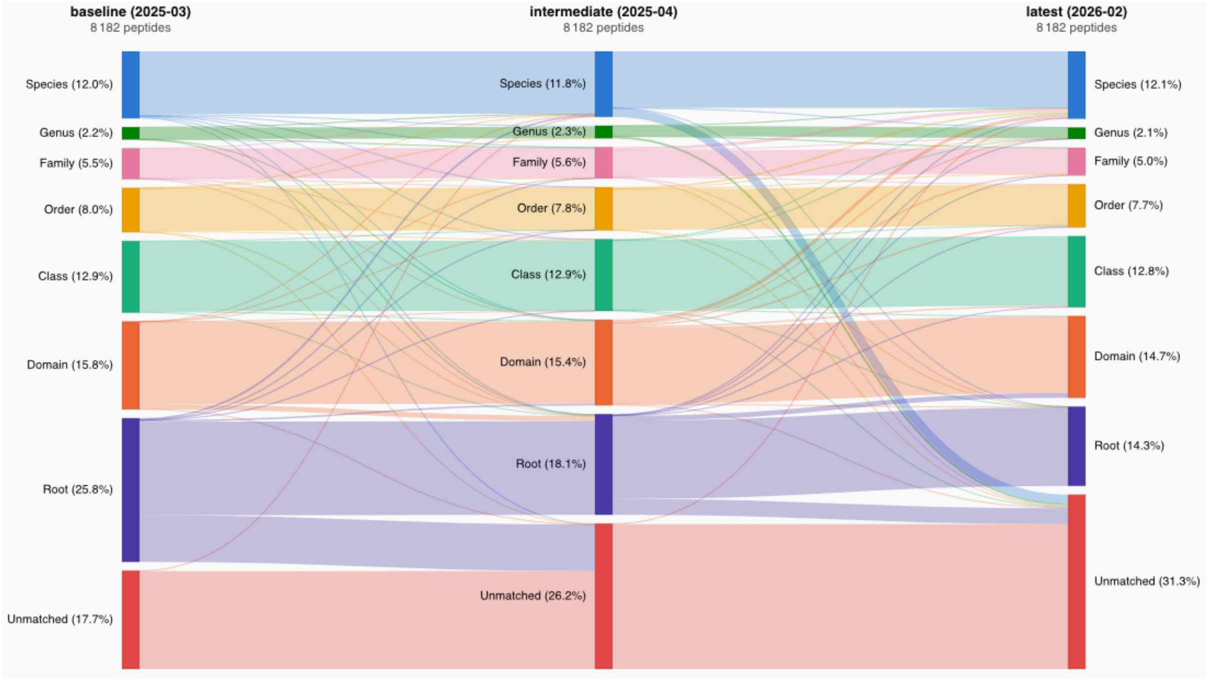


**Figure S1:** Sankey diagram showing peptide-level transitions among exact LCA-rank categories for the marine hatchery dataset across three successive UniProtKB releases. The three columns correspond, from left to right, to UniProtKB 2025_03 (baseline), 2025_04 (intermediate), and 2026_02 (latest). Flows between adjacent columns show how each peptide's exact LCA changed from one release to the next. Each node is a mutually exclusive exact-LCA category (root, domain, class, order, family, genus, species, or unmatched). Node values are shares of all 8 182 input unique peptides and sum to 100% only when the unmatched category is included, so they are not cumulative.

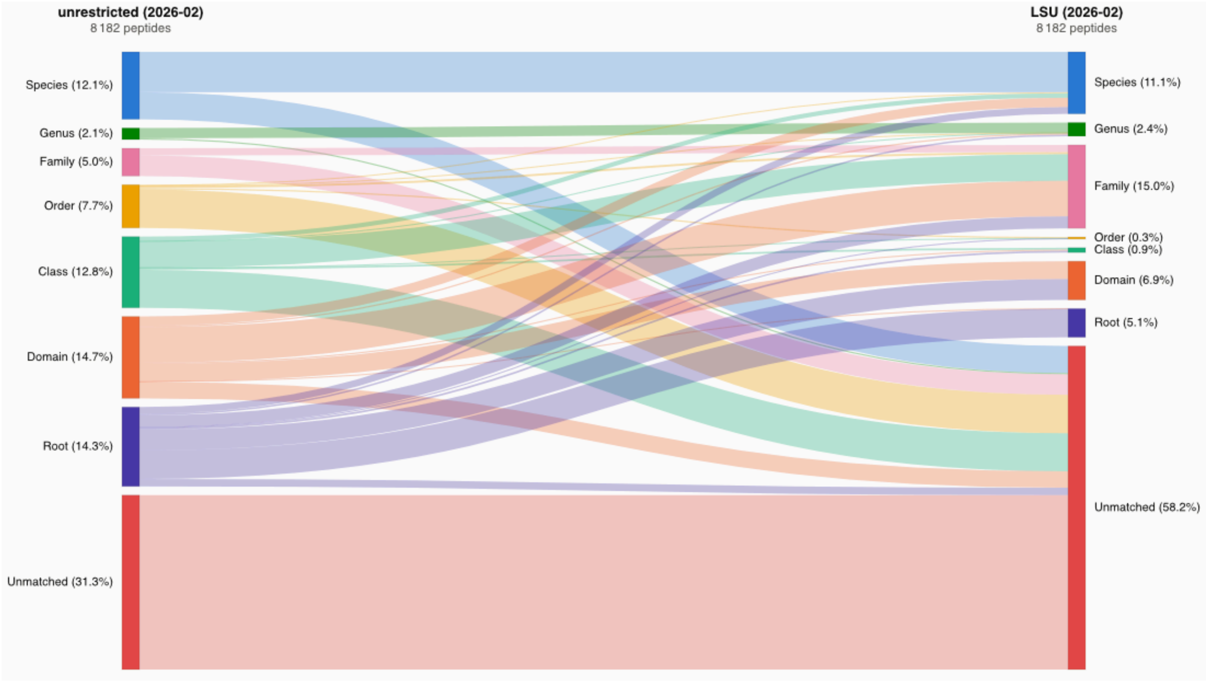


**Figure S2:** Sankey diagram showing marine peptide assignments between two UniProtKB 2026_02 configurations. The two columns correspond, from left to right, to the unrestricted 2026_02 release (unrestricted) and the LSU-restricted database (LSU). Flows between the columns show how each peptide's exact LCA changed under restriction. Each node is a mutually exclusive exact-LCA category (root, domain, class, order, family, genus, species, or unmatched). Node values are shares of all 8 182 input unique peptides and sum to 100% only when the unmatched category is included, so they are not cumulative.
